# Purification of recombinant α-Synuclein: a comparison of commonly used protocols

**DOI:** 10.1101/2020.05.13.093286

**Authors:** Amberley D. Stephens, Dijana Matak-Vinkovic, Gabriele S. Kaminski Schierle

**Affiliations:** Chemical Engineering and Biotechnology, University of Cambridge, Philippa Fawcett Drive, Cambridge, CB3 0AS; Department of Chemistry, University of Cambridge, Lensfield Road, Cambridge, CB2 1EW

**Keywords:** amyloid, hydrophobic interaction chromatography, native mass spectrometry, intrinsically disordered protein

## Abstract

The insoluble aggregated form of the protein alpha-synuclein (aSyn) is associated with synucleinopathies, such as Parkinson’s Disease, therefore great effort is put into understanding why and how this initially soluble protein misfolds. The initial state of aSyn, e.g. presence of contaminants, adducts, oligomers or degradation products, can greatly influence the outcome of an assay, such as determining its aggregation kinetics. Here, we compare four commonly used protocols for the isolation of recombinant aSyn from *E. coli* by boiling, acid precipitation, ammonium sulphate precipitation and periplasmic lysis followed by ion exchange chromatography and gel filtration. We identified, using non-denaturing electrospray ionisation mass spectrometry of the differently extracted aSyn samples, that aSyn isolated by acid precipitation and periplasmic lysis yielded the highest percentage of monomer, 100% and 96.5% respectively. aSyn purity was again highest in samples isolated by acid precipitation and periplasmic lysis, yet aggregation assays displayed differences in the aggregation rate of aSyn isolated by all four methods.

**Highlights:**

- A rapid protocol; expression day one, two step purification day two.
- The periplasmic lysis-based protocol yielded 95% pure aSyn.
- Acid precipitation and periplasmic lysis-based protocols yielded the highest proportion of monomeric aSyn at 100% and 96.5%, respectively.

## Introduction

The protein alpha-synuclein (aSyn) is the predominant protein found in insoluble aggregates called Lewy bodies and Lewy neurites in the neurons of patients suffering from synucleinopathies, such as Parkinson’s Disease. Major emphasis has been put into studying how this usually soluble and intrinsically disordered protein (IDP) misfolds into highly structured β-sheet containing fibrils. Many studies have used purified recombinant aSyn to investigate its misfolding and cellular toxicity in both in vitro and in vivo experiments. However, there are currently several different recombinant aSyn purification protocols in use in the literature, many of which lack validation and quality control of the recombinant protein used.

As an IDP, it is possible to separate aSyn from other proteins using methods to precipitate structured proteins. By boiling a sample, many proteins undergo heat denaturation where internal bonds become broken, disrupting the structure of structured/globular proteins and leading to precipitation, yet heating leaves IDPs in solution due to their lack of structure. Acid shock works in a similar way by disrupting intramolecular bonds but by altering the charge of the amino acids which leads to altered bonding in the proteins and subsequent precipitation. Ammonium sulphate ((NH_4_)_2_SO_4_) precipitation is also used to purify aSyn. Increasing the percentage of (NH_4_)_2_SO_4_ in solution leads to precipitation of different proteins at different concentrations of (NH_4_)_2_SO_4_ due to favouring hydrophobic interactions and self-association^1^. Periplasmic lysis of aSyn from *E. coli* is a more recent protocol that has been chosen as it is less ‘harsh’ on the protein compared to heating or acid shock. aSyn is naturally trafficked to the periplasm when expressed in *E. coli*. To release aSyn, but not the whole cell contents, the periplasm is lysed through osmotic stress ^2^.

aSyn resides as a dynamic ensemble of conformations in its soluble form and is therefore very sensitive to its surrounding environment^3,4^. It is currently unclear whether the methods employed to purify aSyn can actually influence the conformations formed within the dynamic ensemble or can skew the ensemble and whether they can affect which aggregation-prone or non-aggregation-prone pathways aSyn monomers will take. A few studies have investigated the effect of some isolation treatments on the final recombinant aSyn, two studies investigating the effect of heating on the structure of aSyn showed the presence of C-terminus truncated species after heating the protein to 95-100°C^5,6^. Yet, they reported no differences in the overall structure of full length aSyn by farultraviolet circular dichroism (UV-CD) and nuclear magnetic resonance (NMR) spectroscopy^5^. A purification method using acidification to precipitate unwanted proteins was the favoured method by Giehm, et al.,^6^ and was shown to give a higher purity of aSyn compared to boiling^7^. However, it has been observed that acidification leads to C-terminus charge collapse and alteration of long-range interactions within the aSyn monomer, although again this was observed to be reversible^8,9^. It is currently not clear whether ‘reversible’ changes to the conformation of aSyn actually do disrupt intramolecular bonding and lead to small shifts in the dynamic ensemble of the aSyn conformations, which are not detected by averaging measurements such as NMR or CD, or whether they can influence aggregation rates and/or fibril polymorphism. We have previously shown, using the highly sensitive technique of hydrogen-deuterium exchange mass spectrometry (HDX-MS), that the method of storage does impact the monomeric aSyn structure. Lyophilisation, a commonly used storage method for aSyn, leads to a compaction of aSyn monomers in comparison to freezing, even when reconstituted in buffer. The compaction was not detected in methods such as dynamic light scattering^10^. Lyophilisation leads to the formation of lyophilisation-induced oligomers, which were different in structure to those in the frozen aSyn sample, and an increase in variability during ThT-based aggregation assays. Therefore, the treatment of aSyn prior to experiments could be crucial in terms of interpreting experimental data.

Here, we present a comparison of four aSyn isolation methods, boiling, acid precipitation (ppt), (NH_4_)_2_SO_4_ ppt and periplasmic lysis followed by ion exchange chromatography and gel filtration to investigate the purity, proportion of monomer, aggregation rate and fibril polymorphs of aSyn formed.

## Methods and Materials

### *E. coli* expression of recombinant aSyn

The plasmid pT7-7 containing human aSyn cDNA was transformed into *Escherichia coli* One Shot^®^ BL21 (DE3) Star™ (Thermo Fisher Scientific, USA). 0.5 L cultures of *E. coli* in Lysogeny Broth (LB) containing carbenicillin (100 μg/mL) were grown at 37°C with shaking at 200 rpm and induced for expression of aSyn when the OD_600_ reached 0.6-0.8 with 1 mM isopropyl-β-thiogalactopyranoside (IPTG). After four hours of aSyn expression the cells were pelleted by centrifugation at 8 k x g for 15 mins.

### Preparation of protein samples for chromatography

#### Precipitation

For acid ppt and (NH_4_)_2_SO_4_ ppt methods, first the *E. coli* pellet from 500 mL of culture were resuspended in 50 mL of lysis buffer (10 mM Tris, 1 mM EDTA pH 7.2 with protease inhibitor tablets (cOmplete™, EDTA-free protease inhibitor cocktail, Merck, UK)) and sonicated 30 s on and 30 s off for three rounds using a XL-2020 sonicator (Heat Systems, USA). The sonicates were centrifuged at 20 k x g for 30 min and the supernatant saved.

#### Acid precipitation

the pH of the supernatant was reduced to pH 3.5 using HCl and stirred at room temperature (RT) for 20 mins, then centrifuged at 60 k x g for 30 minutes. The pH of the supernatant was then brought up to pH 7.5 with NaOH and stored overnight at 4 °C ^11^.

#### (NH_4_)_2_SO_4_ precipitation

47% w/vol of (NH_4_)_2_SO_4_ was added to the supernatant^12^ and stirred at RT for 20 mins, then centrifuged at 60 k x g for 30 minutes. The pellet of protein was resuspended in 60 mL dialysis buffer (10 mM Tris, 1 mM EDTA pH 7.5) and dialysed overnight against the same buffer at 4 °C.

#### Boiling

For isolation of aSyn by boiling the *E. coli* pellet from 0.5 L of culture was resuspended in 50 mL high salt buffer (0.75 M NaCl, 100 mM MES, 1 mM EDTA pH 7)^13^ and boiled in a water bath for 20 minutes at 100°C, then centrifuged at 60 k x g for 30 minutes. The supernatant was dialysed against 10 mM Tris 1 mM EDTA pH 7.5 overnight at 4 °C in SnakeSkin™ dialysis tubing, with a molecular weight cut off (MWCO) of 10 kDa (Thermo Fisher Scientific, USA).

#### Periplasmic lysis

aSyn was released from the *E. coli* periplasm by osmotic lysis^2^. The pellet of *E. coli* from 0.5 L of culture which had not been frozen was resuspended in 100 mL of osmotic shock buffer (30 mM Tris, 40% sucrose (w/vol), 2 mM EDTA pH 7.2) and incubated at RT for 10 minutes. The solution was centrifuged at 18 k x g for 20 minutes. The supernatant was discarded, and the pellet resuspended in 90 mL of ice cold dH_2_O with 37.5 μL of saturated MgCl_2_ and kept on ice for 3 minutes before centrifuging at 18 k x g for 20 minutes. The supernatant was dialysed overnight against 10 mM Tris, 1 mM EDTA pH 7.5 at 4 °C. The use of EDTA is particularly important after the addition of MgCl_2_ as it influences structure and aggregation rates^14^.

### Ion exchange chromatography

All buffers and aSyn samples for chromatography were filtered through a 0.22 μm filter and degassed before use. For ion exchange chromatography (IEX) the protein was loaded onto a HiPrep Q FF 16/10 anion exchange column (GE Healthcare, Sweden) and eluted against a linear gradient of 7 column volumes (CV) of IEX buffer B (10 mM Tris, 0.75 M NaCl, pH 7.5) followed by 2 CV of 100% IEX buffer B using an ÄKTA Pure fast protein liquid chromatography (FPLC) system (GE Healthcare). To determine the point of elution of aSyn from the chromatography column protein fractions which were collected and monitored on absorption at 280 nm were ran on a 4-12% Bis-Tris gel (Invitrogen, Thermo Fisher) using SDS-PAGE and stained with Coomassie blue. Fractions containing protein bands corresponding to the predicted monomer aSyn molecular weight (MW) of 14.4 kDa were further used in chromatographic steps. Fractions containing aSyn were pooled together and either dialysed overnight in 20 mM Tris pH 7.2 and concentrated with a 10 K MWCO centrifugal concentrator to the desired concentration, ~130 – 140 μM and stored at −80 °C or were directly concentrated before gel filtration.

### Hydrophobic interaction chromatography

For aSyn isolated from the periplasm, we further optimised the purification protocol to yield a higher purity of aSyn by the addition of a hydrophobic interaction chromatography (HIC) step. We changed the counter ion from 0.75 M NaCl to 0.15 M (NH_4_)_2_SO_4_ during IEX chromatography to remove a dialysis step needed to exchange salts before subsequent HIC. The amount of (NH_4_)_2_SO_4_ in the aSyn solution post IEX was calculated based on the % of Buffer B it eluted at. Based on the volume of aSyn protein collected, the amount of (NH_4_)_2_SO_4_ needed to make the solution up to 1M was calculated and then gradually added while stirring.

For HIC, the pH of the aSyn sample was adjusted to pH 7 and filtered through a 0.22 μm filter before being loaded onto a HiPrep Phenyl FF 16/10 (High Sub) column (GE Healthcare, Sweden) and eluted in HIC buffer A (50 mM Bis-Tris, 1 M (NH_4_)_2_SO_4_ pH 7) against a linear gradient of 7 CV of HIC buffer B (50 mM Bis-Tris pH 7) followed by 2 CV of 100% IEX buffer B. Fractions containing aSyn were pooled and extensively dialysed against 20 mM Tris pH 7.2 overnight at 4 °C. The protein solution was concentrated in 10 K MWCO centrifugal concentrators to the desired concentration, ~130 – 140 μM, and frozen at −80 °C until further use.

### Gel filtration

An aliquot of aSyn was defrosted and 500 μL of aSyn was injected into a gel filtration (GF) column, Superdex 75 10/300 GL (GE Healthcare). The sample was eluted by isocratic elution at 0.8 mL/min in 20 mM Tris pH 7.2. Tubing between the injection point and the fraction collector on the ÄKTA Pure FPLC system was changed from orange (0.5 mm) to blue (0.25 mm) to reduce dilution of the protein sample and give a narrower collection peak. Monomeric aSyn eluted at ~ 9 mL.

### Densitometry to determine purity of aSyn

Fractions of proteins samples were run on 4-12% Bis-Tris gels using SDS-PAGE for separation of proteins based on size. The gels were stained with Coomassie blue and the gel image analysed using ImageJ software^15^ to determine the percentage of aSyn present. Regions of interest were selected and a histogram of the intensity of dyed protein in the area displayed. From the histogram the area of aSyn was calculated as a percentage of the total area of stained proteins to give the percentage purity.

### Reverse phase high pressure liquid chromatography to determine the purity of aSyn

The purity of the aSyn samples was analysed by analytical reversed phase chromatography (aRP) on a 1260 Infinity high pressure liquid chromatography (HPLC) system (Agilent Technologies LDA UK Limited, UK), equipped with an autosampler and a diode-array detector. 50 μL of sample was injected onto a Discovery BIO Wide Pore C18 column (15 cm x 4.6 mm, 5 μm column with a guard column) (Supelco, Merck, UK) and eluted on a gradient of 95% water + 0.1% acetic acid and 5% acetonitrile + 0.1% acetic acid to 5% water + 0.1% acetic acid and 95% acetonitrile + 0.1% acetic acid at a 0.8 mL/min flow-rate over 40 mins. The elution profile was monitored by UV absorption at 220 and 280 nm. The area under the peaks in the chromatograph of absorption at 280 nm was calculated to provide the percentage purity of aSyn. aSyn eluted at ~17.9 mins.

### Native mass spectrometry

Non–denaturing nano-electrospray ionization mass spectrometry (Native mass spectrometry) was used to analyse the oligomerisation states of recombinant aSyn prepared in four different ways: boiled, (NH_4_)_2_SO_4_ ppt, acid ppt and periplasmic lysis. Native mass spectra were recorded on a Synapt HD mass spectrometer (Waters, Manchester, UK) modified for studying high masses. Protein samples were exchanged into 0.20 M ammonium acetate (pH 7.0) solution using Micro Bio–Spin 6 chromatography columns (Bio–Rad, USA) and diluted to a final concentration of 5–10 μM before analysis. An aliquot of 2.5 μL of protein solution was electrosprayed from a borosilicate emitter (Thermo Scientific, UK) for sampling. Typical conditions for the data acquire were capillary voltage 1.6-2.2 kV, cone voltage 160–190 V, Trap 40-50 V, Transfer 140 V with backing pressure 3–4 mbar and source temperature of 20 °C. Spectra were calibrated externally using caesium iodide. Data acquisition and processing were performed using MassLynx 4.1. Spectra were edited manually using Adobe Illustrator for the purpose of this publication.

### Thioflavin-T based kinetic aggregation assays

20 μM freshly made thioflavin-T (ThT) (abcam, Cambridge, UK) was added to 50 μL of 20 μM aSyn after GF in 140 mM KCl, 20 mM Tris pH 7.2. All samples were loaded onto nonbinding, clear bottom, 96-well half-area plates (Greiner Bio-One GmbH, Germany). The plates were sealed with a SILVERseal aluminium microplate sealer (Grenier Bio-One GmbH). Fluorescence measurements were taken using a FLUOstar Omega plate reader (BMG LABTECH GmbH, Ortenbery, Germany). The plates were incubated at 37°C with double orbital shaking at 300 rpm for five minutes before each read every hour for 170 hours. Excitation was set at 440 nm with 20 flashes and the ThT fluorescence intensity measured at 480 nm emission with a 1300 gain setting. ThT assays were repeated twice using four wells for each condition. For aSyn isolated by boiling, four gel filtrated samples were used, for aSyn isolated by (NH_4_)_2_SO_4_ ppt three gel filtrated samples were used, for aSyn isolated by acid ppt two gel filtrated samples were used, and for aSyn isolated by periplasmic lysis four gel filtrated sampled were used. Data were normalised to the sample with the maximum fluorescence intensity for each plate.

### Analytical size exclusion chromatography to determine the remaining aSyn monomer concentration after aggregation assays

At the end of the ThT-based aggregation assays, the amount of remaining monomer of aSyn in each well was determined by analytical size exclusion chromatography on a HPLC (SEC-HPLC). The contents of each well after the ThT-based assay were centrifuged at 21k x g for 20 minutes and the supernatant was added to individual aliquots in the autosampler of the Agilent 1260 Infinity HPLC system (Agilent Technologies LDA UK Limited, UK). 25 μL of each sample was injected onto an Advance Bio SEC column, 7.8 x 300 mm 300Å (Agilent, UK) in 20 mM Tris pH 7.2 at 1 mL/min flowrate. Injections were also made for each sample at the start of the assay to quantify the amount of starting protein. The elution profile was monitored by UV absorption at 220 and 280 nm. The remaining monomer percentage was calculated from the ratio of the area under the curves at the beginning and the end of the assay.

### Transmission Electron Microscopy

20 μL of aSyn was taken directly from the ThT-based aggregation assay plates of the boiled, (NH_4_)_2_SO_4_ ppt and periplasmic lysis samples and diluted 1:4 with dH_2_O. The acid ppt aSyn sample was used neat, and all samples were incubated on glow-discharged carbon coated copper grids for 1 min before washing twice with dH_2_O. 2% uranyl acetate was used to negatively stain the samples for 30 s before imaging on the Tecnai G2 80-200kv transmission electron microscopy (TEM) at the Cambridge Advanced Imaging Centre.

## Results

### Acid precipitation of *E. coli* proteins leads to the highest purity of aSyn before chromatography

Four commonly used protocols for the purification of recombinant aSyn were compared to determine which yielded the highest proportion of monomeric aSyn and the highest degree of purity. First, in all four protocols, 0.5 L of *E. coli* culture was induced for four hours with IPTG before pelleting the bacteria. The pellets were then treated differently dependent on the isolation protocol. For boiled samples, the *E. coli* pellet was resuspended in a high salt buffer then boiled in a water bath at 100 °C for 20 minutes before centrifuging. The supernatant was dialysed overnight in 10 mM Tris, 1 mM EDTA pH 7.5. For the acid and (NH_4_)_2_SO_4_ ppt protocols, the *E. coli* pellets were resuspended in 10 mM Tris, 1 mM EDTA pH 7.5 including protease inhibitors before being sonicated and centrifuged. The supernatant was then precipitated either by reducing the pH to 3.5 with HCl on a stirrer for 20 mins or by addition of 47% (w/vol) (NH_4_)_2_SO_4_ on a stirrer for 20 mins. The precipitates were centrifuged, the supernatant from the acid ppt was brought back to neutral pH with NaOH and stored overnight at 4 °C before chromatography was performed. The pellet from the (NH_4_)_2_SO_4_ ppt containing the aSyn was resuspended in 10 mM Tris, 1 mM EDTA pH 7.5 and dialysed overnight. For periplasmic lysis, the *E. coli* periplasm was lysed by osmotic shock. The *E. coli* pellet was resuspended in a sucrose based buffer which acts as an osmotic stabiliser preventing whole cell lysis^16^. After centrifugation to pellet the *E. coli*, the outer membrane was lysed by osmotic shock with water and MgCl_2_ to release the contents of the periplasm, but not the cytoplasm. The lysed protein was dialysed overnight in 10 mM Tris, 1 mM EDTA pH 7.5. The acid precipitated aSyn sample was the most pure at this point, with 96.5% purity by densitometry measurement of the SDS-PAGE Coomassie stained gel, which agrees with a previous study^7^ (Supplementary Figure 1, Supplementary Table 1).

### Chromatographic isolation of aSyn yields 80-95% pure aSyn

IEX was then used to isolate aSyn from all protein solutions using a HiPrep™ Q FF 16/10 anion exchange column. aSyn was eluted on a linear gradient of IEX buffer A (10 mM Tris, 1 mM EDTA pH 7.5) against IEX buffer B (10 mM Tris, 1 mM EDTA, 0.75 M NaCl, pH 7.5) (Figure 1 a.i, b.i, c.i, d.i). To determine which fractions the aSyn resided in, the samples were analysed by SDS-PAGE and the gel stained by Coomassie blue to visualise the protein. Fractions containing aSyn are highlighted in the coloured block on the IEX chromatograms (Figure 1 a.i,ii, b.i,ii, c.i,ii, d.i,ii). The purity of the samples was analysed by densitometry and the aSyn precipitated in acid and the aSyn that was boiled were found to be 100% and 99.3% pure, respectively (Supplementary Table 1). After IEX the aSyn samples were dialysed in 20 mM Tris pH 7.2 and concentrated using centrifugal concentrators with a MWCO of 10 kDa. aSyn was concentrated until the protein concentration was between 130-140 μM before storage at −80°C. To increase the purity of the aSyn from samples isolated by periplasmic lysis and (NH_4_)_2_SO_4_ ppt further, and to ensure isolation of monomeric protein, gel filtration was used. 500 μL of aSyn was injected onto a Superdex 75 10/300 GL column and eluted isocratically (Figure 1, a.iii, b.iii, c.iii, d.iii.) The Coomassie blue stained gel after SDS-PAGE of aSyn showed that the all isolation protocols, apart from periplasmic lysis (Figure 1d.iv), lead to 100% pure aSyn after IEX and GF (Figure 2a., Supplementary Table 1). The aSyn sample isolated by periplasmic lysis still contained contaminating proteins, aSyn was only 91.7% pure in fraction 1 and 97.7% pure in fraction 2 (Figure 1div stars indicate the contaminants, Supplementary Table 1). Previous protocols using the periplasmic lysis protocol have also employed an extra hydrophobic interaction chromatography (HIC) step^17^. An additional HIC step was added using a HiPrep™ Phenyl Fast Flow (high sub) 16/10 column, but the previous protocol was updated to save time by substituting the counter ion salt in IEX from NaCl to (NH_4_)_2_SO_4_ to prevent an additional buffer exchange step before HIC. Therefore, directly after IEX (NH_4_)_2_SO_4_ was added to make the protein solution up to 1 M (NH_4_)_2_SO_4_, equivalent to the starting buffer A for HIC. aSyn was eluted on a linear gradient against HIC buffer B (50 mM Bis Tris, pH 7) (Figure 1d.v, d.vi). aSyn was 100% pure when analysed by densitometry after IEX, HIC and GF (Figure 2a., Supplementary Table 1). Supplementary Figure 2 and Supplementary Table 2 shows a second purification run for each purification method and the concentration of protein in each step of purification, showing the methods to be reproducible.

**Figure 1.**
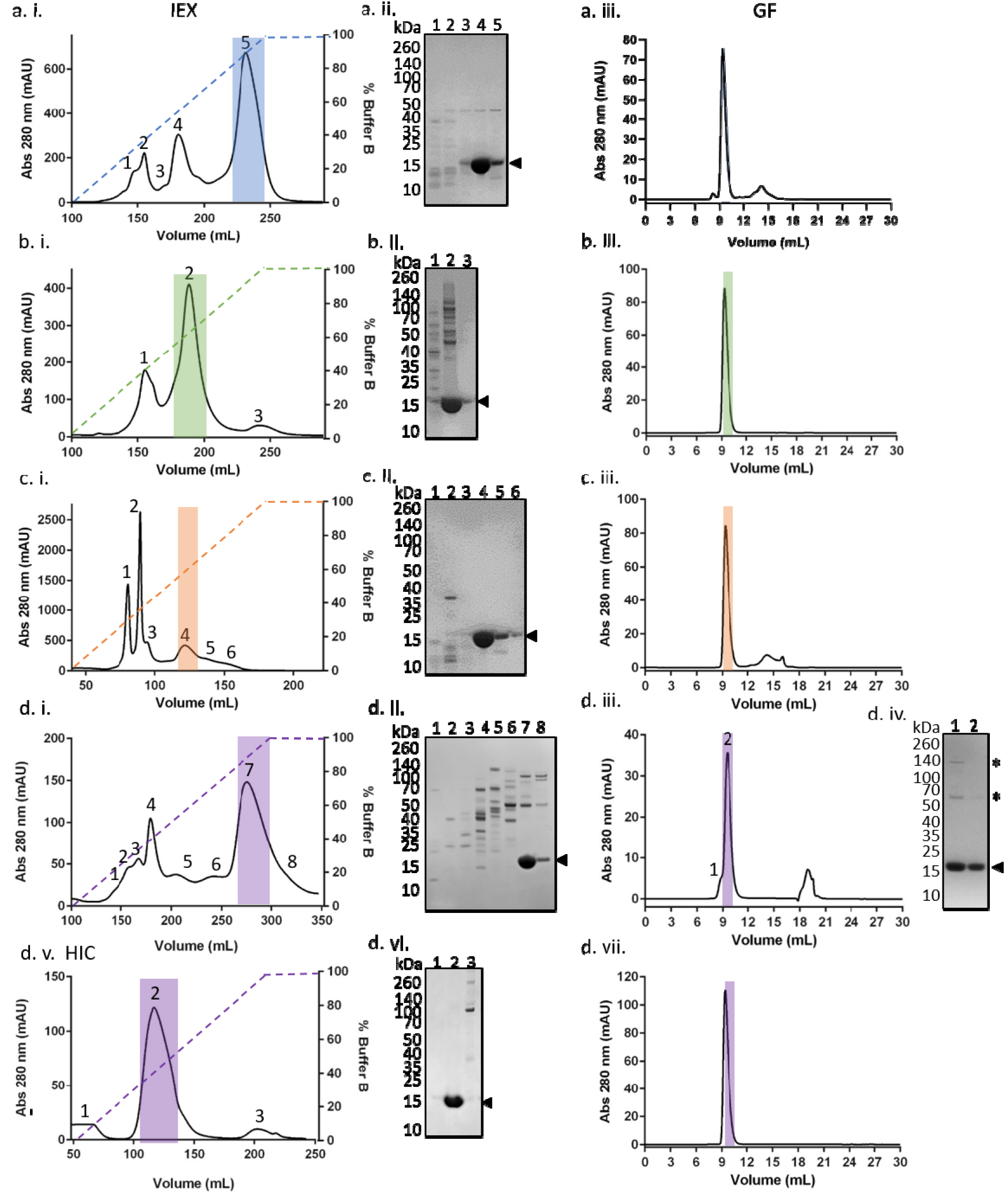
aSyn isolated by boiling, acid ppt and (NH_4_)_2_SO_4_ ppt is highly pure after IEX and GF, while aSyn isolated by periplasmic lysis requires an additional HIC step to increase purity. aSyn was isolated by IEX from samples that were (a.i.) boiled, (b.i.) precipitated by (NH_4_)_2_SO_4_, (c.i.) precipitated by acidification and (d.i.) lysed from the periplasm. Protein fractions from IEX were taken from individual peaks with maximum absorption at 280 nm and analysed by SDS-PAGE using a 4-12% bis-tris gel which was stained by Coomassie blue, (a.ii.) boiled, (b.ii.) precipitated by (NH_4_)_2_SO_4_, (c.ii.) precipitated by acidification and (d.ii.) lysed from the periplasm. aSyn ran at ~ 15 kDa, indicated by the arrow next to the gel images, and the peak aSyn resided in is highlighted in colour on the IEX chromatographs. GF of pooled fractions containing aSyn after IEX show monomeric aSyn eluting after ~9 mL, (a.iii.) boiled, (b.iii.) precipitated by (NH_4_)_2_SO_4_, (c.iii.) precipitated by acidification and (d.iii.) periplasmic lysis. GF of aSyn isolated by periplasmic lysis did not yield as highly pure aSyn as the other methods did, (d.iv.) as indicated by the presence of contaminating proteins (*) in the Coomassie blue stained gel. Therefore, an additional (d.v.) HIC step was added and (d.vi.) aSyn was shown to be subsequently purer as shown by the Coomassie blue stained gel. (d.vii.) The final GF of aSyn isolated by periplasmic lysis after IEX and HIC showed a single peak of monomeric aSyn eluting at ~9 mL.

**Figure 2.**
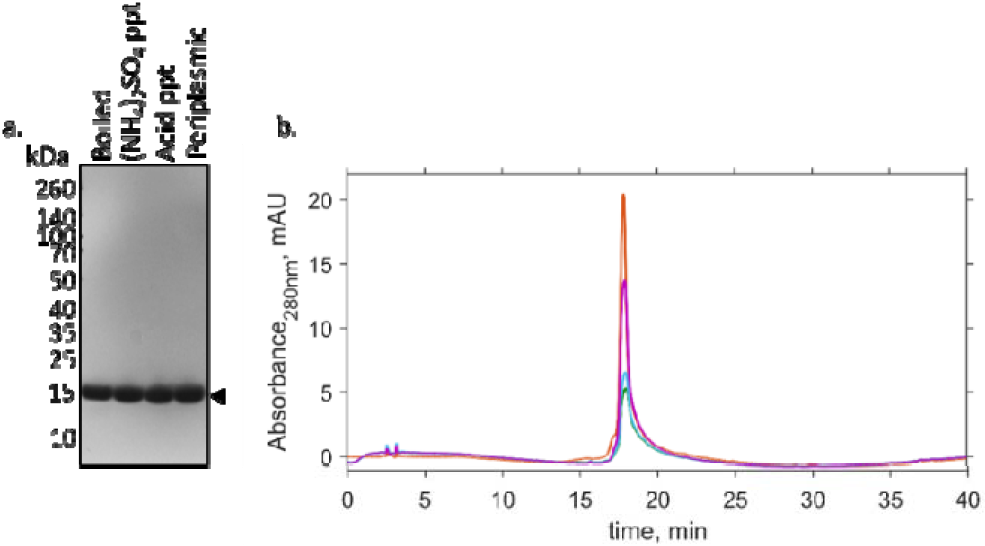
Coomassie blue stained gel and reverse phase chromatographs of aSyn after gel filtration show highly pure monomeric aSyn. (a.) Samples of aSyn after gel filtration were analysed by SDS-PAGE on a 4-12% Bis-Tris gel and stained with Coomassie blue. aSyn appears as a single band, indicated by the arrow at around 15 kDa. (b.) 50 μL of each sample was injected onto an analytical Discovery BIO Wide Pore C18 column to determine the purity of aSyn. aSyn isolated by boiling (blue) was 86% pure, aSyn isolated by (NH_4_)_2_SO_4_ ppt (green) was 81% pure, aSyn isolated by acid ppt was (orange) 89.9% pure and aSyn isolated from periplasmic lysis (purple) was 95% pure, determined by the area under the peak.

Densitometry analysis of the Coomassie blue stained gel of the four aSyn samples after GF showed 100% pure monomeric aSyn in all samples. However, analytical reversed phase chromatography (aRP) was also employed to determine the purity of each sample as it is a more sensitive method to detect contaminants (Figure 2b). The samples were shown to be less pure after IEX by aRP compared to densitometry measurements (Supplementary Figure 3a), sample purity ranged from 62.9 % to 84.8% when analysed by aRP, but ranged between 49.7 and 100% pure when analysed by densitometry (Table 1 and Supplementary Table 1). aRP of aSyn purified by IEX, HIC and GF compared to only IEX and GF led to an increase in purity from 63.5% to 95% (Supplementary Figure 3b). After GF the aSyn purity was determined to be 86% for the boiled sample, 81% for (NH_4_)_2_SO_4_ ppt, 89.9% for acid ppt and 95% for periplasmic lysis of aSyn by aRP (Figure 2b, Table 1).

**Table 1.** Purity of aSyn at different steps of isolation determined by reverse phase chromatography

|  | Boiled |  |  | (NH<sub>4</sub>)<sub>2</sub>SO<sub>4</sub> ppt |  |  | Acid ppt |  |  | Periplasmic lysis |  |  |
| --- | --- | --- | --- | --- | --- | --- | --- | --- | --- | --- | --- | --- |
|  | Total protein (mg/mL) | % purity | Final aSyn conc. (μM) | Total protein (mg/mL) | % purity | Final aSyn conc. (μM) | Total protein (mg/mL) | % purity | Final aSyn conc. (μM) | Total protein (mg/mL) | % purity | Final aSyn conc. (μM) |
| Post IEX | 1.038 | 69.3 |  | 0.619 | 84.8 |  | 1.59 | 62.9 |  | 0.458 | 82.5 |  |
| Post HIC | - | - |  | - | - |  | - | - |  | 0.254 | 86.7 |  |
| GF | 0.219 | 86 | 36.9 | 0.267 | 81 | 44.7 | 0.257 | 89.9 | 42.9 | 0.292 | 95 | 48.7 |

### Analysis of aSyn samples by native mass spectrometry shows that the acid precipitation protocol yields the most highly monomeric aSyn

Obtaining highly pure monomeric aSyn is needed for the majority of assays performed. Although the SDS-PAGE Coomassie blue stained gel and aRP methods show monomeric aSyn, the SDS used in the PAGE and the organic solvents used in aRP are denaturing and may give a false impression of the level of monomeric protein present. Instead, we employed non-denaturing nano electrospray ionization mass spectrometry (native MS). The technique permits the study of protein structure at physiological pH and the identification of aSyn multimers and degradation products without the need to use cross-linkers which may alter structure or induce artefacts^18^. In the boiled sample, aSyn was found in both monomer and dimer form, but also as a degraded product of 11562±3 Da comprising 36.6% of the sample (Figure 3a, Table 2). A degraded product of 12172 Da was also identified by Giehm, et al., after boiling of aSyn^6^. The percentage of aSyn products were calculated by the relative intensity of the m/z peaks in each charge state (Supplementary Table 3). As the degraded product was not detected by SDS-PAGE or aRP it may have been induced during electrospray ionisation. aSyn samples precipitated in (NH_4_)_2_SO_4_ contained monomer (90.3%), dimer (8.5%) and trimer (1.2%) (Figure 3b, Table 2), while the aSyn isolated by acid ppt contained only monomeric aSyn (Figure 3c, Table 2). aSyn isolated by periplasmic lysis was highly monomeric (96.5%) with a small percentage of dimer (3.5%) (Figure 3d, Table 2). The monomer was disordered in all samples, as expected, and the dimer and trimers were possibly linked by non-covalent bonds which remain formed during non-denaturing electrospray ionisation.

**Figure 3.**
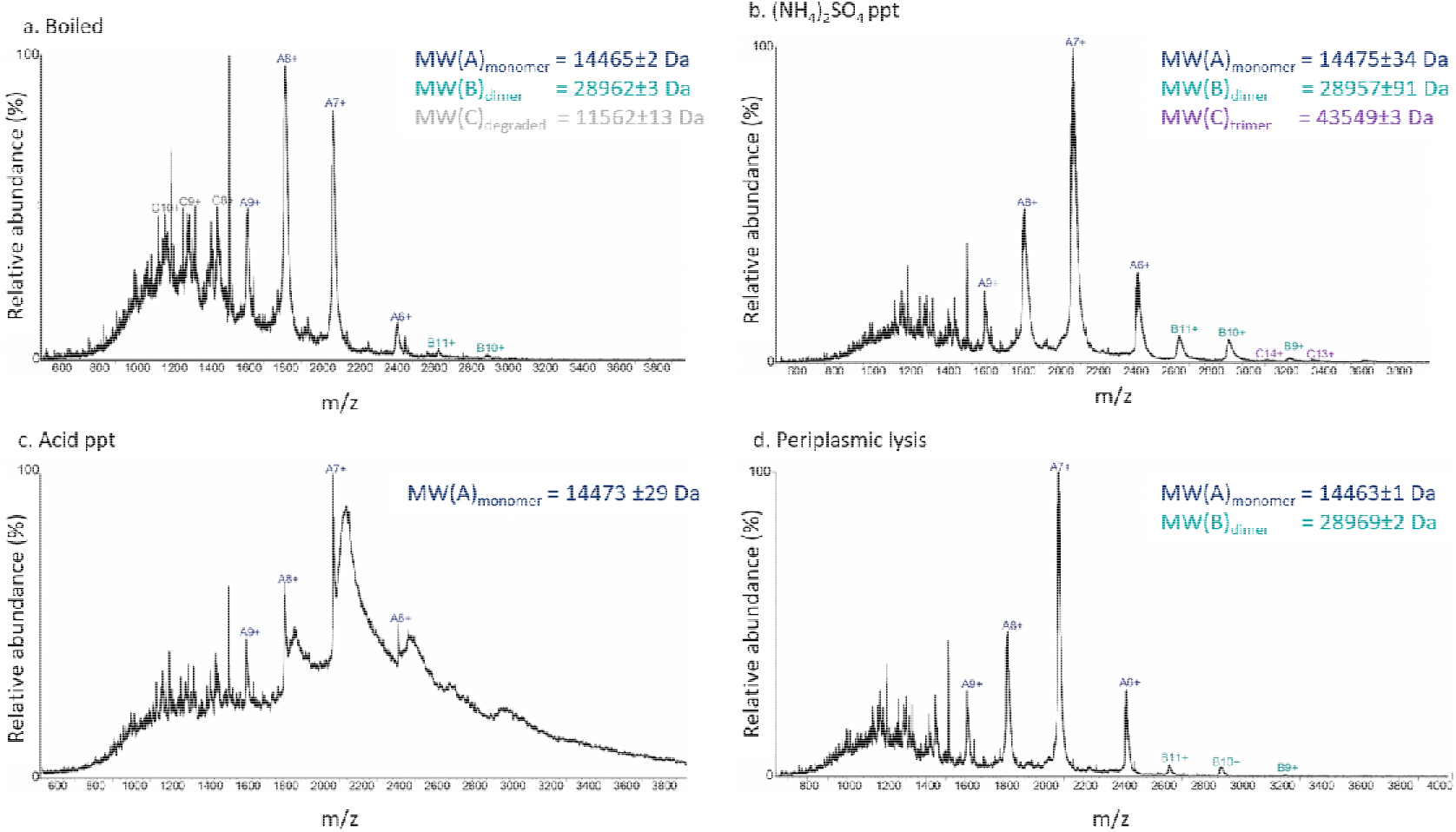
Native MS data of aSyn after gel filtration shows that acid precipitation yields the highest percentage of monomeric aSyn. aSyn in 200 mM NH_4_CH_3_CO_2_ was analysed by native MS. aSyn isolated by (a.) boiling was found as a monomer (A) in charge states 9+ to 6+, the MW is highlighted in blue, as a dimer (B), highlighted in teal, in charge states 10+, 11+ and as a potentially degraded product (C) highlighted in grey at charge states 10+ to 8+ with a MW of ~11562 Da. aSyn isolated with (b.) (NH_4_)_2_SO_4_ ppt was identified as monomeric (A), dimeric (B) and trimeric (C) (~43549 Da) protein forms, the trimer charge states 14+, 13+ and MW are highlighted in purple. The aSyn sample isolated with (c.) acid ppt was only found in a monomeric state (A), while aSyn isolated by (d.) periplasmic lysis was found to be in monomeric (A) and dimeric (B) forms.

**Table 2.** Molecular weight and relative percentage of aSyn structures in each sample determined by native nano-ESI-MS

| | Boiled | | $(\text{NH}_4)_2\text{SO}_4$ ppt | | Acid ppt | | Periplasmic lysis | |
| --- | --- | --- | --- | --- | --- | --- | --- | --- |
|  | MW | % | MW | % | MW | % | MW | % |
| Degraded | 11562 ± 13 | 36.6 | - | - | - | - | - | - |
| Monomer | 14465.4 ± 2.2 | 60.2 | 14475 ± 34 | 90.3 | 14473 ± 29 | 100 | 14463 ± 1 | 96.5 |
| Dimer | 28962.5 ± 3.0 | 3.2 | 28957 ± 91 | 8.5 | - | - | 28969 ± 2 | 3.5 |
| Trimer | - | - | 43549 ± 3 | 1.2 | - | - | - | - |

To determine whether the difference in percentage of monomeric, dimeric and trimeric aSyn affects the aggregation rate, we performed kinetic aggregation assays using the molecule ThT which fluoresces when bound to fibrillated forms of aSyn, providing a kinetic readout^19^. 20 μM of aSyn was incubated with 20 μM of ThT in 20 mM Tris, 100 mM KCl pH 7.2 for one week. The kinetic aggregation curves show that the aSyn isolated by periplasmic lysis was the most aggregation prone, followed by aSyn isolated by (NH_4_)_2_SO_4_ ppt (Figure 4a). Surprisingly, aSyn isolated by boiling and acid ppt appeared to be the least aggregation prone under the conditions tested (Figure 4a). As it is known that ThT assays are highly variable and that ThT also has varying fluorescence intensities when bound to different fibril polymorphs^19,20^, we also performed analytical size exclusion chromatography on a HPLC (SEC-HPLC) to determine the quantity of remaining aSyn monomer. We observe that the quantity of the remaining aSyn monomer does not fully reflect the ThT fluorescence observed, where at least 40% of the acid ppt aSyn appears to have formed higher order structures (Figure 4b). However, the remaining monomer concentration does reflect the trend observed in the ThT-based assays, whereby the aSyn isolated by periplasmic lysis had the least remaining monomer and aSyn isolated by (NH_4_)_2_SO_4_ ppt also had less remaining monomer than the samples that were boiled and precipitated by acid. We further investigated the morphology of the aSyn samples to determine whether fibrils had formed and whether their morphology differed using TEM. TEM showed fibrils present in all samples, but fibrils were harder to find in the sample from acid ppt, indicating less fibrils were present. All fibrils have straight morphology (Figure 4c, Supplementary Figure 4), as shown previously for aSyn aggregated in the presence of salt^21^. The fibril bundles also showed lateral binding (Figure 4c, shown by the arrows).

**Figure 4.**
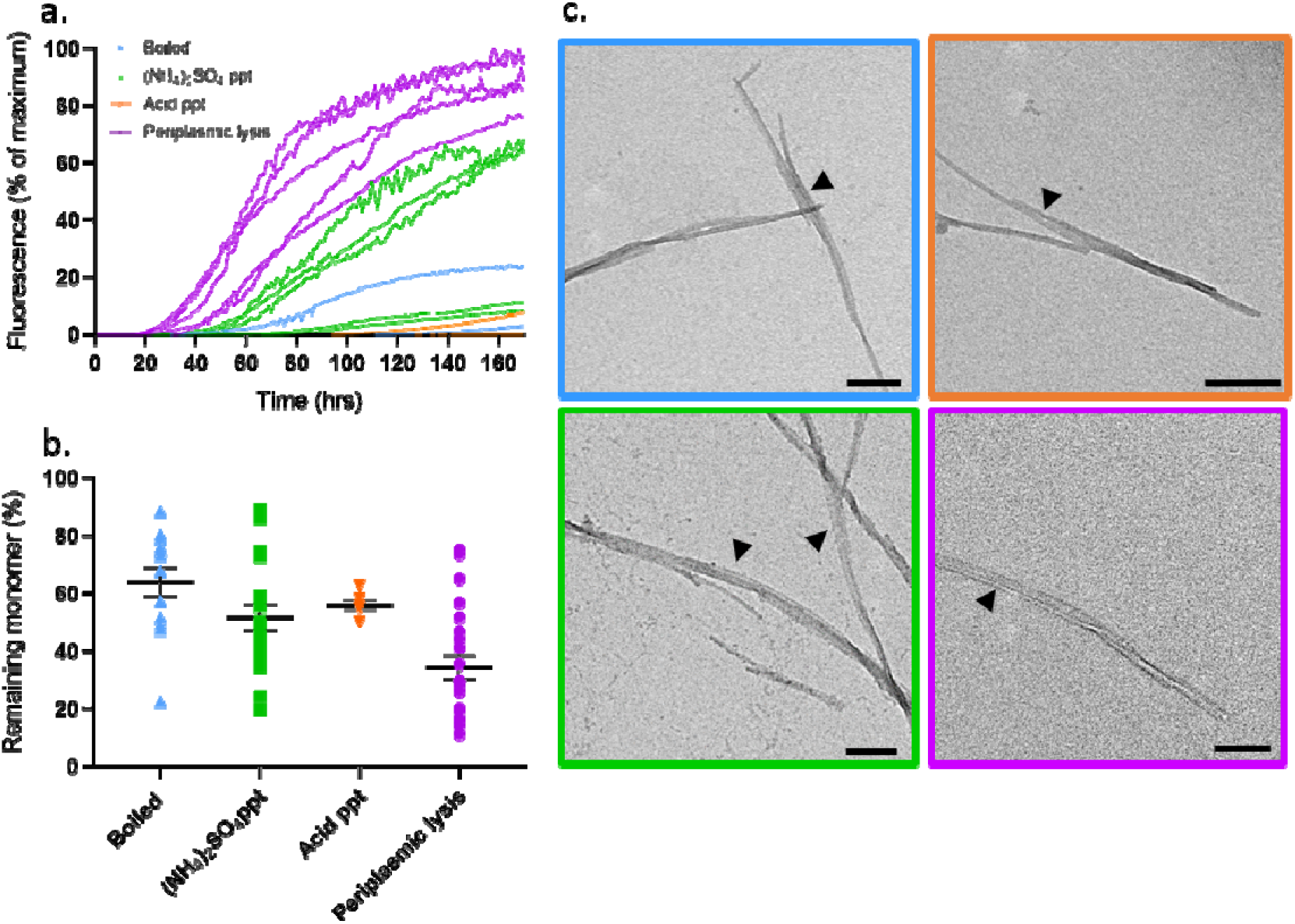
Kinetic aggregation assays show aSyn isolated by periplasmic lysis is the most aggregation prone and all aSyn samples have straight fibril morphology. (a.) ThT-based aggregation assays show aSyn isolated by periplasmic lysis (purple, n=5) and (NH_4_)_2_SO_4_ ppt (green, n=5) aggregate at a faster rate and to a greater extent than aSyn isolated by boiling (blue, n=4) or acid ppt (orange, n=3). Each sample (n) represents aSyn from individual GF runs plated as four well replicates over two plates. 20 μM aSyn in 20 mM Tris, 100 mM KCl pH 7.2 was incubated with 20 μM ThT in a half area 96 well plate with double orbital agitation at 300 rpm for 5 minutes before each read every hour for 170 hours. (b.) The percentage of the remaining monomer concentration in each well after the ThT assay was determined by performing SEC-HPLC and calculating the area under the curve compared to the area under the curve of the starting monomeric sample. Error bars represent SEM from wells n=14 boiled, n=19 (NH_4_)_2_SO_4_, n=7 acid, n=22 periplasmic. (c.) aSyn samples were taken directly from the ThT wells, boiled (blue), (NH_4_)_2_SO_4_ ppt (green) and periplasmic lysis (purple) samples were diluted 1:4 and the acid precipitated (orange) sample was used neat when incubating on grids before being imaged by TEM. All samples showed fibrils with straight morphology and lateral binding between fibril bundles (black arrows). Scale bar = 200 nm.

## Discussion

Many different protocols are currently used for the purification of aSyn, yet little investigation has been performed into the exact product that is present at the end of these purification methods. Here, we compared four commonly applied protocols, boiling, acid ppt, (NH_4_)_2_SO_4_ ppt and periplasmic lysis to determine which of the four methods yielded the highest protein purity and the most monomeric aSyn. Isolation of aSyn by acid ppt and periplasmic lysis yielded the highest percentage of monomeric protein, at 100% and 96.5% respectively, and 89.9% and 95% purity, respectively. The aggregation rate of aSyn prepared by these two methods varied greatly when monitored using a ThT-based kinetic assay, where aSyn isolated by acid ppt barely aggregated, yet aSyn isolated by periplasmic lysis aggregated well under the conditions used. The remaining monomer concentration did not reflect the amount of fibrillisation of aSyn isolated by acid ppt, possibly indicating that more oligomeric structures or amorphous aggregates had formed which were not detected by ThT fluorescence or SEC-HPLC. This may indicate that the method of aSyn isolation can impact the aggregation propensity of the dynamic ensemble of monomer conformations.

Further work is needed to determine whether the purification protocols we use can influence the conformation dynamics of aSyn. Currently techniques are not sensitive enough to determine if we are skewing the dynamic equilibrium of conformations by using different purification techniques, which could then impact the propensity of aSyn to aggregate, or even the subsequent fibril polymorphs formed and its toxicity^22^. It is thus important to characterise the sample fully to guarantee reproducibility and validity of the data.

## Author Information

### Author Contributions

A.D.S Designed experiments and performed purification, kinetic assays and TEM. D.M-V. performed native MS. A.D.S and G.S.K wrote the manuscript. All authors have given final approval of the manuscript.

### Notes

The authors declare no competing financial interests.

## Acknowledgements

We would like to thank Dr Penny Hamyln for helpful discussions on chromatography protocols. Lyn Carter and Filomena Gallo from the Cambridge Advanced Imaging Centre for help with sample loading into the TEM. This work was supported by Wellcome Trust, Alzheimer’s Research UK Grants, the Michael J Fox Foundation and Infinitus China Ltd.

## Supplementary Information

Raw data is available at the University of Cambridge Data Repository.

## Supplementary Information

**Supplementary Figure 1.**
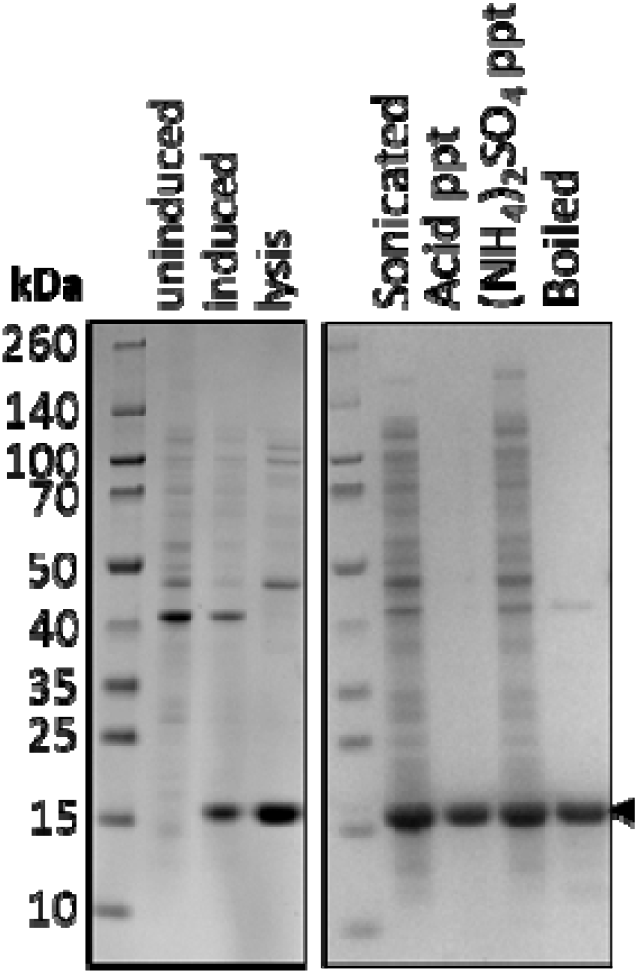
Coomassie blue stained 4-12% Bis-Tris gel of aSyn samples isolated by different methods shows acid precipitation yields the highest purity of aSyn prior to chromatography. Samples of E. coli before (uninduced) and after induction (induced) with 1 mM IPTG show expression of aSyn ~15 kDa shown by the black arrow in the induced lane. aSyn was then extracted from E. coli by different methods: aSyn lysed from the periplasm after osmotic shock is shown as periplasmic lysate (lysis). Prior to precipitation methods, aSyn was sonicated to break open the E. coli (sonicated) before i) acid precipitation and recovery of the supernatant containing aSyn by centrifugation (Acid ppt), ii) precipitation with (NH_4_)_2_SO_4_ and iii) recovery of aSyn from the pellet after centrifugation ((NH_4_)_2_SO_4_ ppt). E. coli were also boiled and the aSyn recovered in the supernatant after centrifuging (Boiled).

**Supplementary Table 1.** Purity of aSyn determined by densitometry of Coomassie blue stained gels

| Isolation protocol | aSyn % purity |  |  |
| --- | --- | --- | --- |
|  | PreIEX | Post IEX | Post GF |
| Boiled | 91.1 | 99.3 | 100.0 |
| $(\text{NH}_4)_2\text{SO}_4$ ppt | 37.6 | 49.7 | 100.0 |
| Acid ppt | 96.5 | 100.0 | 100.0 |
| Periplasmic Lysis | 70.0 | 92.2 | Fr1 91.7<br>Fr2 97.7 |

**Supplementary Figure 2.**
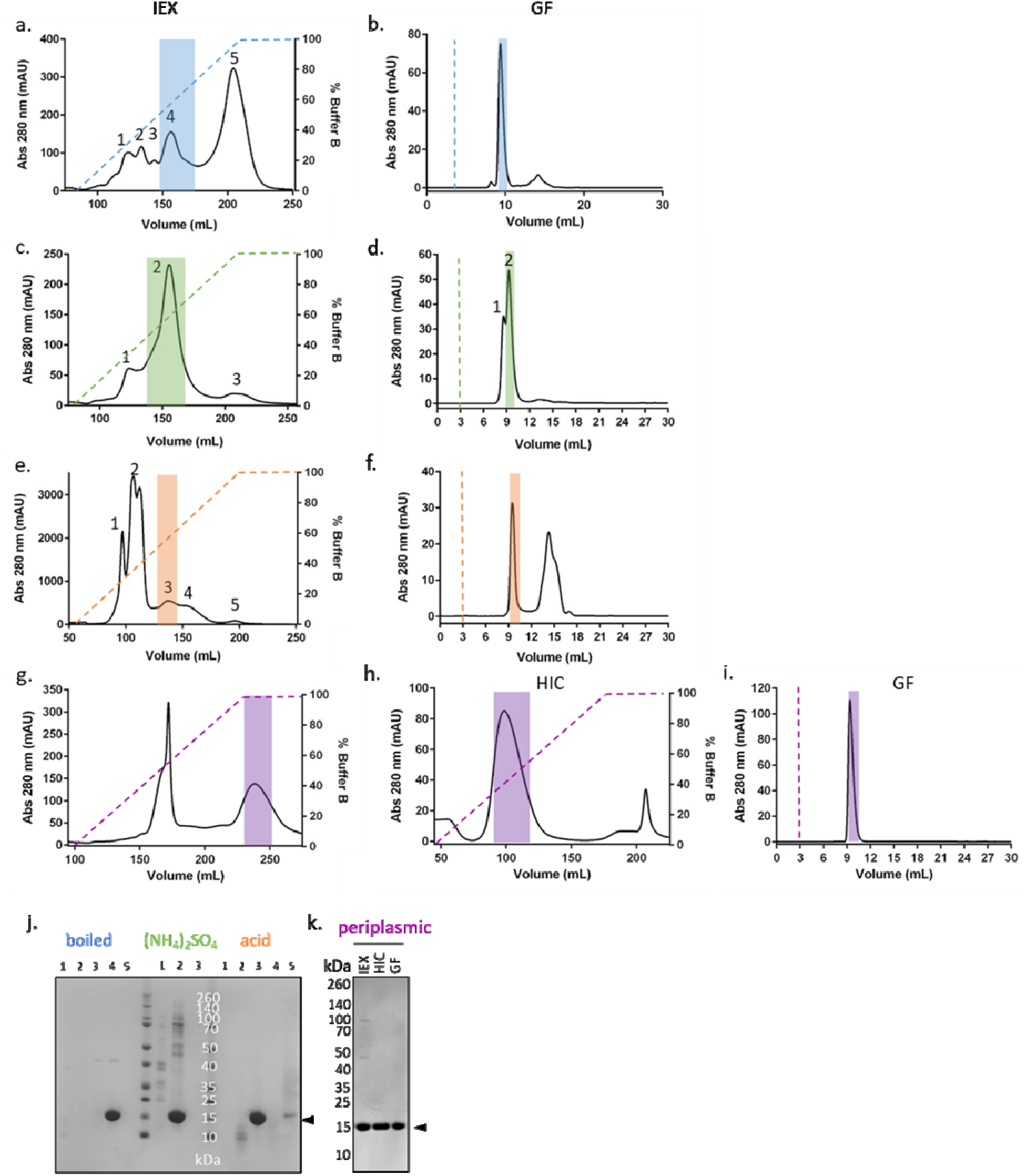
The second purification run also shows that unlike the other isolation protocols aSyn isolated by periplasmic lysis requires an additional HIC step to increase purity. Chromatograph of IEX chromatography of aSyn isolated by (a.) boiling E. coli, (c.) (NH_4_)_2_SO_4_ precipitation, (e.) acid precipitation and (g.) periplasmic lysis, followed by GF for the (b.) boiled aSyn, (d.) (NH_4_)_2_SO_4_ precipitation and (f.) acid precipitation. aSyn from periplasmic lysis was further purified by (h.) hydrophobic interaction chromatography (HIC) before (i.) GF. A Coomassie blue stained gel of fractions separated by SDS-PAGE from the eluted protein peaks of the IEX chromatography were used to determine which peak aSyn resided in (j), shown by the black arrow ~ 15 kDa, and correspond to the highlighted regions in the IEX graphs (a.,c.,e.). (k) Protein fractions from IEX, HIC and GF for aSyn from periplasmic lysis show aSyn by the black arrow and correspond to the highlighted areas in g., h., i.,.

**Supplementary Table 2.**
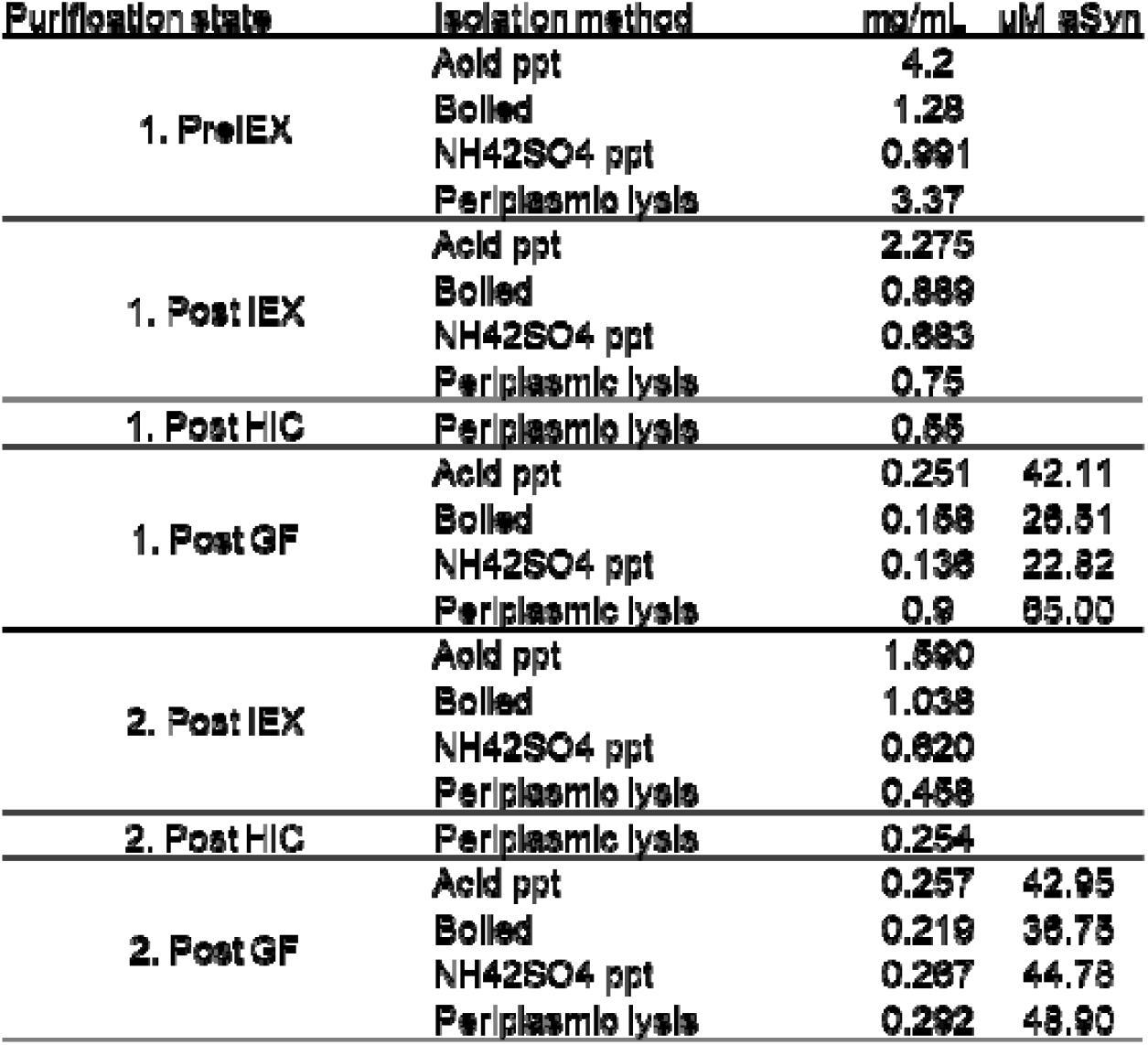
Concentration of total protein and aSyn during purification run one and two

**Supplementary Figure 3.**
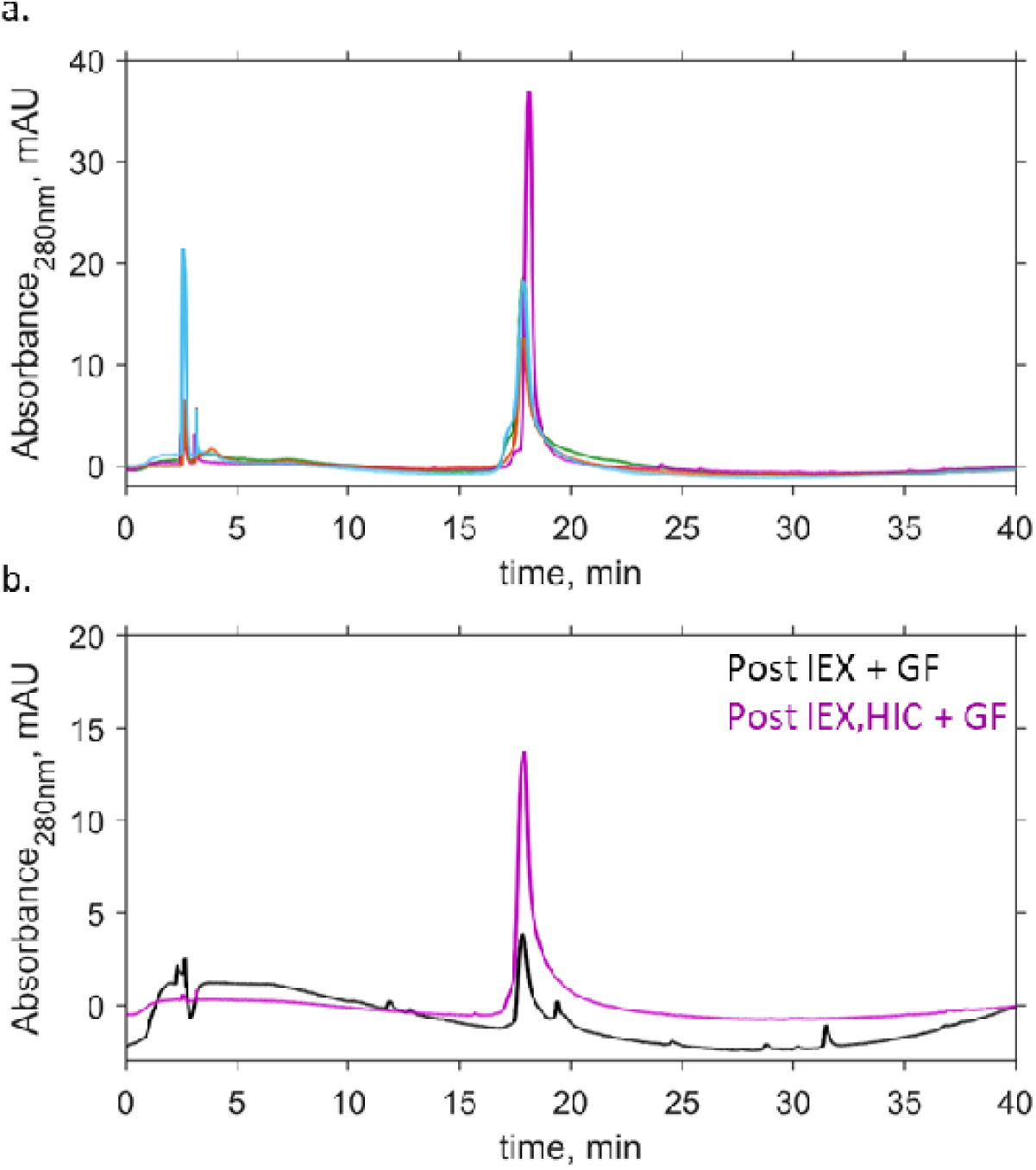
Analytical reverse phase chromatography shows more impurities than Coomassie blue stained gels. Pooled fractions of aSyn after IEX were analysed by analytical reverse phase chromatography (aRP) to determine purity. 50 μL of sample was injected into a C18 column and aSyn was eluted on a gradient of 95% water with 0.1% acetic acid and 5% acetonitrile with 0.1% acetic acid at 0.8 mL/min. (a) The purity of aSyn isolated by boiling (blue), acid precipitation (orange), (NH_4_)_2_SO_4_ precipitation (green) and periplasmic lysis (purple) was determined from the area under the peaks, aSyn eluted ~ 17.8 mins. (b) Purity of aSyn after IEX and GF (black) compared to IEX, HIC and GF (purple) shows fewer contaminating proteins in the sample which had the additional HIC step.

**Supplementary Table 3.**
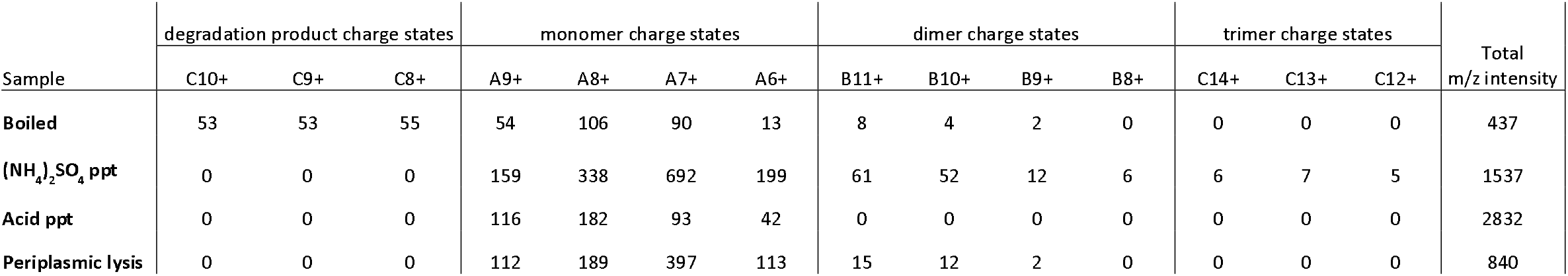
Peak intensity of degraded protein, monomer, dimer and trimers from native MS used to calculate the percentage of aSyn structures in each sample

**Supplementary Figure 4.**
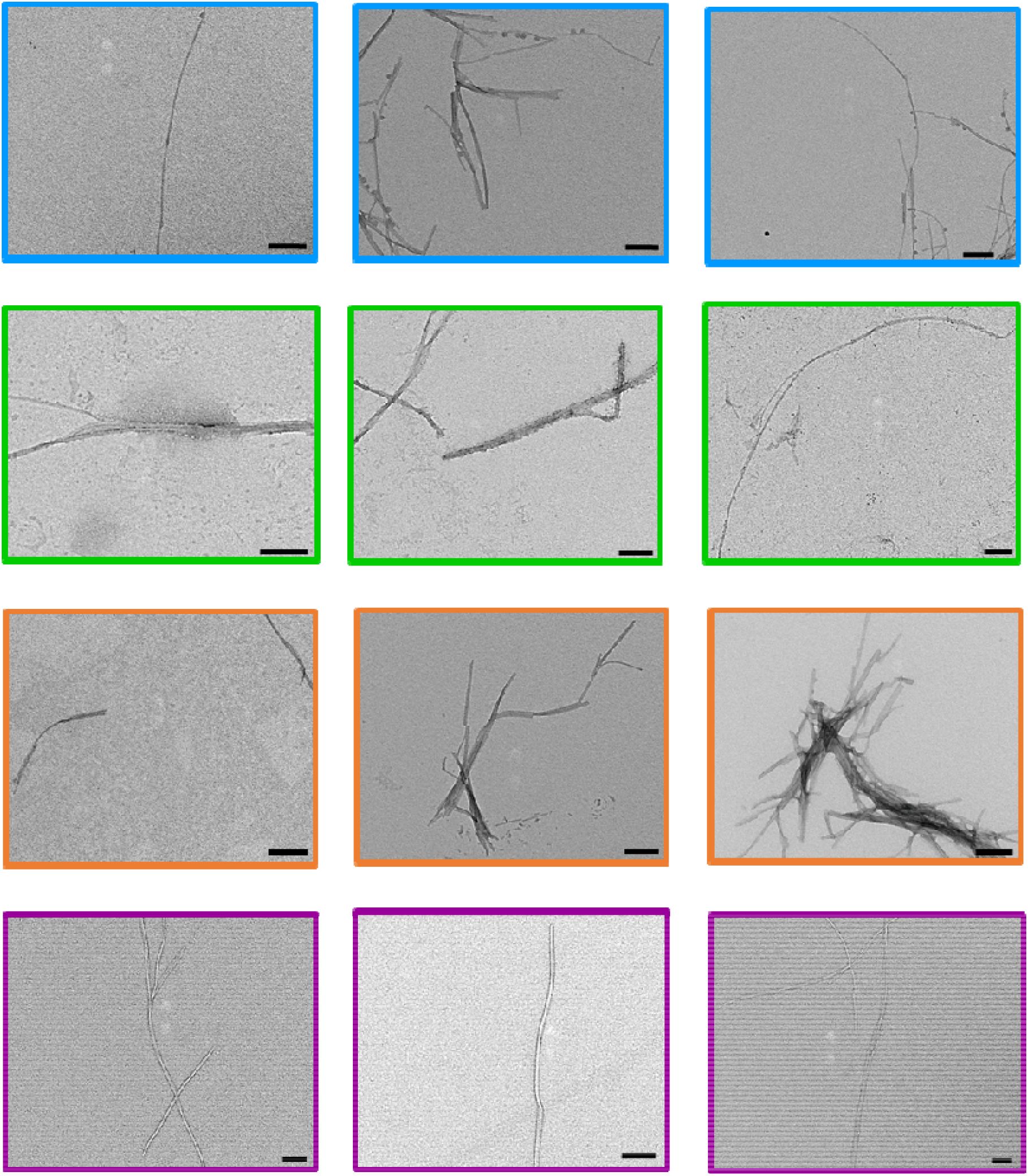
Straight aSyn fibrils are formed during ThT assays from all purification methods used. aSyn samples from the ThT assays were imaged by TEM and show a straight fibril morphology. The aSyn taken from wells of sample isolated by boiling (blue), (NH_4_)_2_SO_4_ precipitation (orange) and periplasmic lysis (purple) were diluted to 5 μM before imaging. The aSyn isolated by acid precipitation (orange) was incubated on the grid directly from the well at 20 μM. Representative images are shown. Scale bar = 200 nm.

**Supplementary Figure 5.**
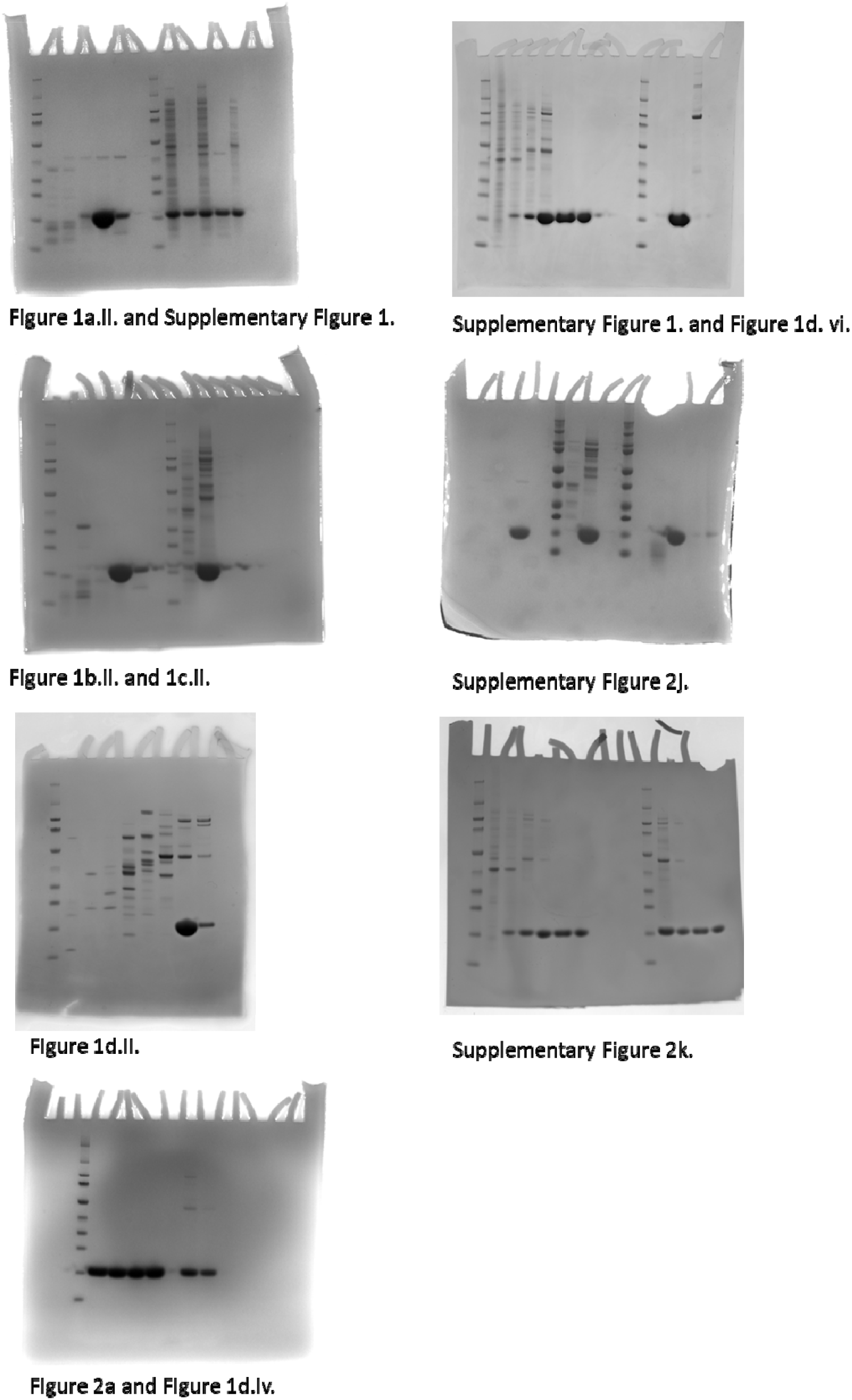
All raw images of Coomassie blue stained SDS-PAGE gels used in this publication.

